# *Cirrina*: LLM-driven pharmacological reasoning agent enables preclinical CNS drug evaluation

**DOI:** 10.64898/2026.03.29.713781

**Authors:** Binita Rajbanshi, Khalid Iqbal, Anuj Guruacharya

## Abstract

Assessing whether a preclinical drug candidate will work is not a prediction problem but a reasoning problem. The same numerical output warrants different interpretations depending on the target and therapeutic context. CNS drug development presents the most demanding instance of this reasoning problem. For example, a compound must cross the blood–brain barrier, resist efflux transport, and achieve adequate receptor occupancy at a dose that clears safety margins. The constraints interact with each other in a web that needs careful interpretation. Here, we show that Cirrina, an LLM agent coupled to eight mechanistic pharmacology tools, can reason across the input data to provide better decisions and a well documented reasoning trace. The LLM agent reasons across multiple data tiers from SMILES to animal PK/PD measurements adjusting thresholds based on target-specific requirements. Validated against 181 CNS compounds, it achieved a 68% accuracy compared to a rule-based deterministic pipeline of 31% accuracy. In 103 discordant cases, the agent’s reasoning was correct in 75% of instances compared to only 10% for deterministic pipelines. Cirrina provides a scalable, documented framework for preclinical decision-making, effectively identifying failure-prone candidates that generic thresholds overlook, and thereby reducing the chances of failure along the clinical development cycle.

## Introduction

CNS drug development presents a particularly demanding instance of this reasoning problem, carrying the highest attrition rate of any therapeutic area, with over 90% of candidates failing between Phase I and approval [1,2]. A CNS drug must cross the blood-brain barrier (BBB) in sufficient quantities, resist efflux transport, achieve adequate receptor occupancy at its brain target, and maintain these properties within a tolerable dose range [3,4]. These requirements are interdependent. Efflux transport determines unbound brain concentration, which determines receptor occupancy, which determines therapeutic dose, which feeds back to safety margins [5,6]. A compound that crosses the BBB adequately may still fail because P-glycoprotein (P-gp) efflux depletes free drugs in the interstitial fluid below the concentration required for target engagement [7]. Conversely, a compound with low predicted brain exposure may succeed because its target requires only 15-20% receptor occupancy rather than the 60-80% typical of dopamine D2 antagonists [8,9]. These outcomes cannot be determined from any single endpoint prediction. They emerge from the interaction between predictions across the pharmacokinetic-pharmacodynamic (PK/PD) chain.

Current computational tools occupy two extremes that both fail to address this integration problem. At one end, freely available ADMET predictors such as pkCSM [10], SwissADME [11], and ADMETlab [12] provide rapid binary classifications of individual endpoints, BBB permeability, P-gp substrate likelihood, hERG inhibition, but treat each property in isolation. They do not model the dependencies between outputs, and the result is a collection of independent Pass/Fail verdicts that cannot explain which mechanism limits a compound. Pharmacologists must mentally assemble these isolated predictions into a coherent assessment, a process that is inconsistent and undocumented. At the other extreme, enterprise physiologically-based pharmacokinetic (PBPK) platforms such as Simcyp [13] and GastroPlus [14] can model mechanistic interactions with physiological fidelity [15,16]. However, these platforms require extensive parameterization, measured hepatic clearance, plasma protein binding, permeability coefficients, transporter kinetics, that is typically unavailable during lead optimization, the stage where compound prioritization decisions are most consequential [17]. Furthermore, enterprise platforms offer no adaptive workflow that adjusts model complexity to data availability.

This creates an integration gap at the centre of computational preclinical assessment. The step that determines translational success, chaining interdependent predictions, interpreting which mechanism is rate-limiting, propagating prediction uncertainty, and applying target-specific pharmacological knowledge is performed manually by experienced pharmacologists. The quality of this integration varies with individual expertise, is rarely documented, and cannot be reproduced [18].

Recent advances in large language models (LLMs) have demonstrated that these systems can function as autonomous agents capable of multi-step reasoning over structured tools [19,20]. In scientific domains, LLM agents have been applied to chemical synthesis planning [21], scalable chemical design [22], molecular property prediction [23], and literature-guided hypothesis generation [24,25]. However, the application of LLM agents to pharmacological reasoning where the agent must chain mechanistic models, interpret intermediate outputs, and integrate target-specific knowledge remains unexplored. Pharmacological assessment is not a prediction problem but a reasoning problem. The same numerical output (e.g., 25% predicted receptor occupancy) warrants different interpretations depending on the target, the confidence in upstream predictions, and the therapeutic context.

Here we present *Cirrina*, an LLM agent that operates as a pharmacological reasoning loop over a mechanistic computational engine. The engine implements eight pharmacology tools, BBB classification, P-gp and BCRP efflux scoring, active transport detection, PK parameter estimation, four-compartment CNS PBPK simulation with saturable efflux, Hill equation receptor occupancy, quantitative safety margin analysis, and inverse PK/PD dose estimation that collectively span the prediction chain from molecular structure to clinical dose. Model complexity adapts to available data through five input tiers, from structure-only (Tier 1) through measured toxicity data (Tier 5). The LLM agent executes tools, examines outputs, decides what to run next based on intermediate results, and re-interprets downstream predictions in light of upstream findings. A parallel deterministic pipeline executing the same engine with fixed decision rules serves as an ablation control, isolating what pharmacological reasoning contributes beyond mechanical integration alone. Architectural schematic of Agentic vs. deterministic frameworks is shown in **Fig 1**.

**Fig 1.**
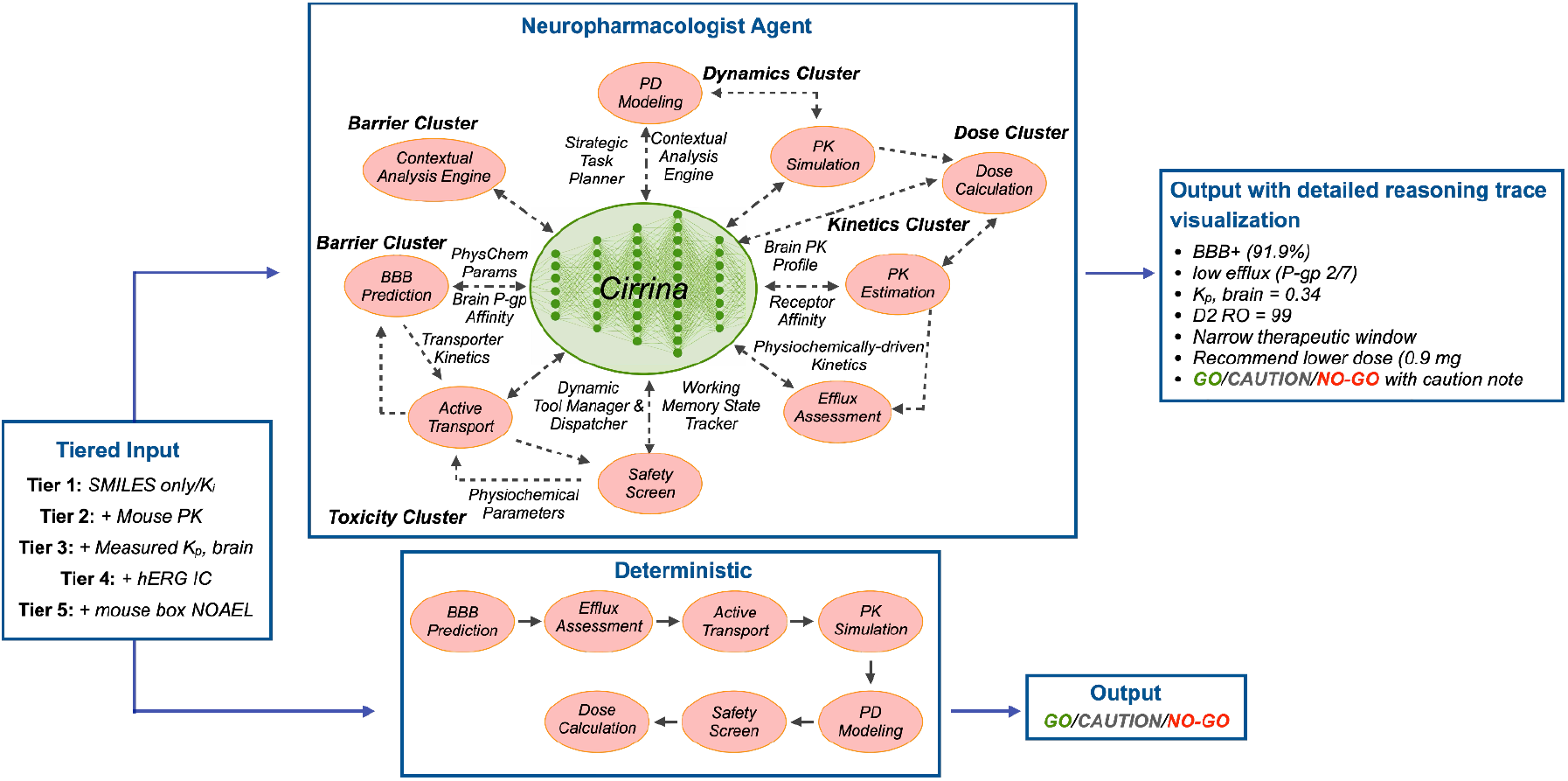
Architectural Schematic of Agentic vs. Deterministic Frameworks. Both architectures process data through a five-tier input structure. The Agentic framework utilizes an LLM orchestrator to manage eight specialized tools, while the Deterministic framework relies on a singular baseline tool. Both systems generate decision outputs categorized as GO, CAUTION, or NO-GO. Notably, the Agentic framework provides granular reasoning traces to justify its decision-making, a feature absent in the deterministic pipeline.

We validate *Cirrina* across 181 CNS compounds spanning 14 molecular targets, demonstrating that LLM-driven pharmacological reasoning over mechanistic tool outputs produces more accurate compound assessments than the same tools with fixed decision rules. The agent’s advantage is concentrated where target-specific pharmacological knowledge overrides generic thresholds precisely the cases where a human pharmacologist would also outperform a fixed rule.

## Results

We evaluated *Cirrina* across 181 CNS compounds comprising 103 validated CNS drugs, 25 externally validated CNS drugs, 32 BBB-negative controls, and 21 NO-GO compounds (16 clinically failed CNS candidates and 5 withdrawn drugs). All compounds were assessed at Tier 1 (structure-only); a subset was additionally evaluated at Tiers 2 and 3 with measured preclinical PK and brain partitioning data. A deterministic pipeline executing the same eight computational tools with fixed decision rules served as the ablation control. The underlying mechanistic engine allometric scaling, blood–brain barrier prediction, receptor occupancy, safety models, and PK conformal intervals are individually validated in **Figs. 2a-2f**. End-to-end uncertainty propagation and sensitivity analyses are reported in **Figs. 2g-2h**.

**Fig 2.**
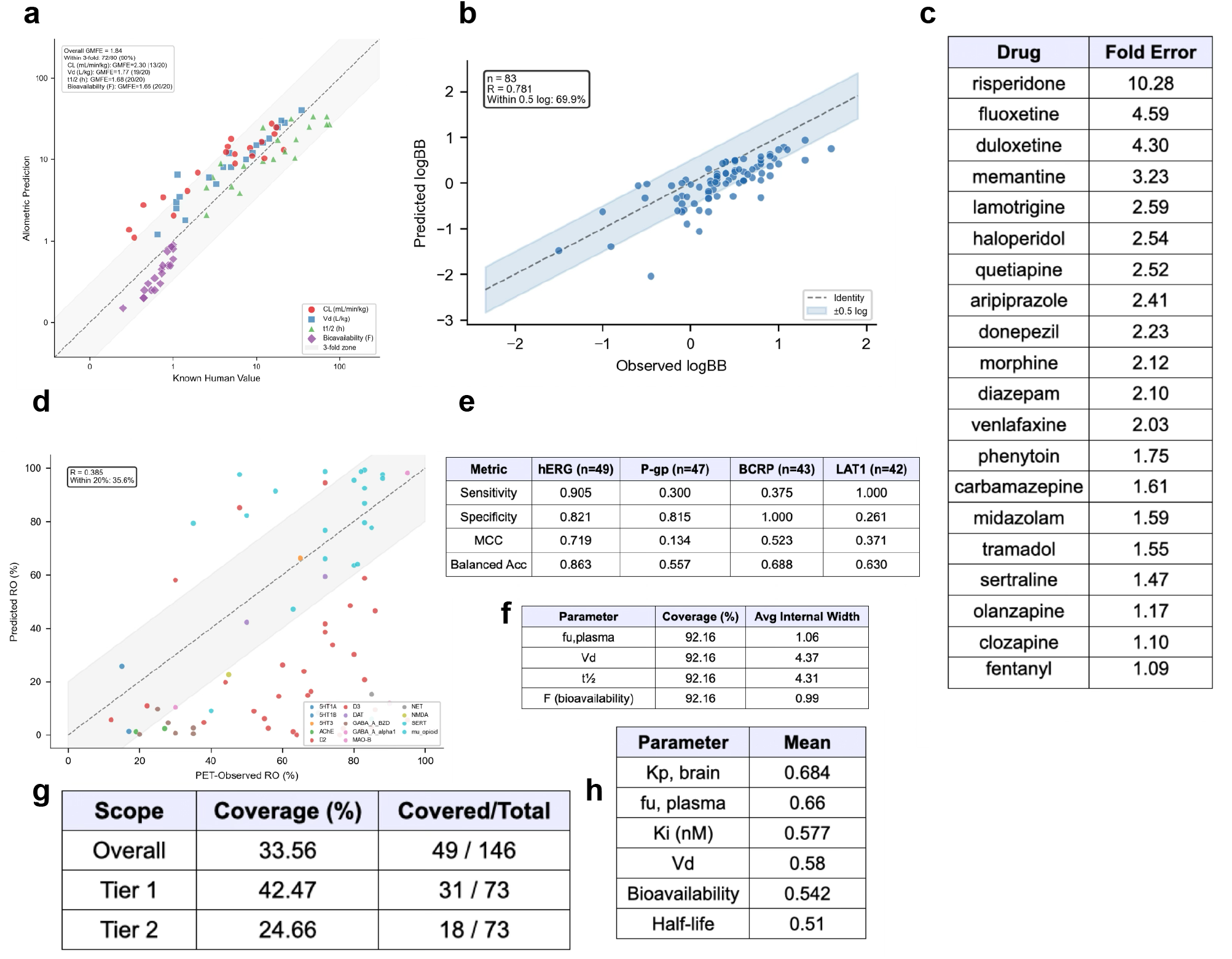
Computational engine validation. **(a)** Allometric scaling validation for PK parameters (n = 20). Predictions within 3-fold error: CL 75%, Vd 95%, t1/2 100%, F 100%. **(b)** Predicted vs. observed logBB correlation (R = 0.781); 69.9% within 0.5 log units. **(c)** Kp,brain fold-error analysis (GMFE = 2.21); 80% within 3-fold. Outliers (risperidone, fluoxetine, duloxetine) likely reflect transporter interactions. **(d)** Receptor occupancy validation across 73 comparisons and 14 targets (R = 0.385); 35.6% within 20% absolute error. **(e)** Safety and transporter prediction balanced accuracy: hERG 0.86, P-gp 0.56, BCRP 0.69, LAT1 0.63. **(f)** Conformal PK coverage of 92.2% at 90% nominal confidence, with interval widths ranging 1.0–4.4x. **(g)** End-to-end uncertainty propagation coverage of 33.6%, well below the 90% target, reflecting compounding uncertainty across multi-tool reasoning. **(h)** Parameter sensitivity ranking: Kp,brain (0.684) and fu,plasma (0.660) are the dominant drivers, followed by Ki and Vd (0.577), F (0.542), and t1/2 (0.51).

### The agent identifies viable and failed CNS compounds

Across all 181 compounds with ground-truth recommendation labels (121 GO, 17 CAUTION, 43 NO-GO), the agent achieved overall recommendation accuracy of 68.0% (123/181) **(Fig. 3a)** compared with 30.9% (56/181) for the deterministic pipeline **(Fig. 3b)**. Because the evaluation set is class-imbalanced (66.9% GO), raw accuracy alone understates the agent’s discriminative ability; class-adjusted metrics reveal a larger separation. The agent achieved Matthews correlation coefficient (MCC) of 0.469 versus 0.075 for the deterministic pipeline (near-random), and AUC-ROC of 0.874 versus 0.577 indicating that the agent ranks compounds well even when hard classification boundaries are imperfect. Critically, the agent detected 76.7% (33/43) of NO-GO compounds compared with 23.3% (10/43) for the deterministic pipeline, and achieved Precision at 10 of 1.00 (the ten highest-risk-ranked compounds were all true NO-GO or CAUTION).

**Fig 3.**
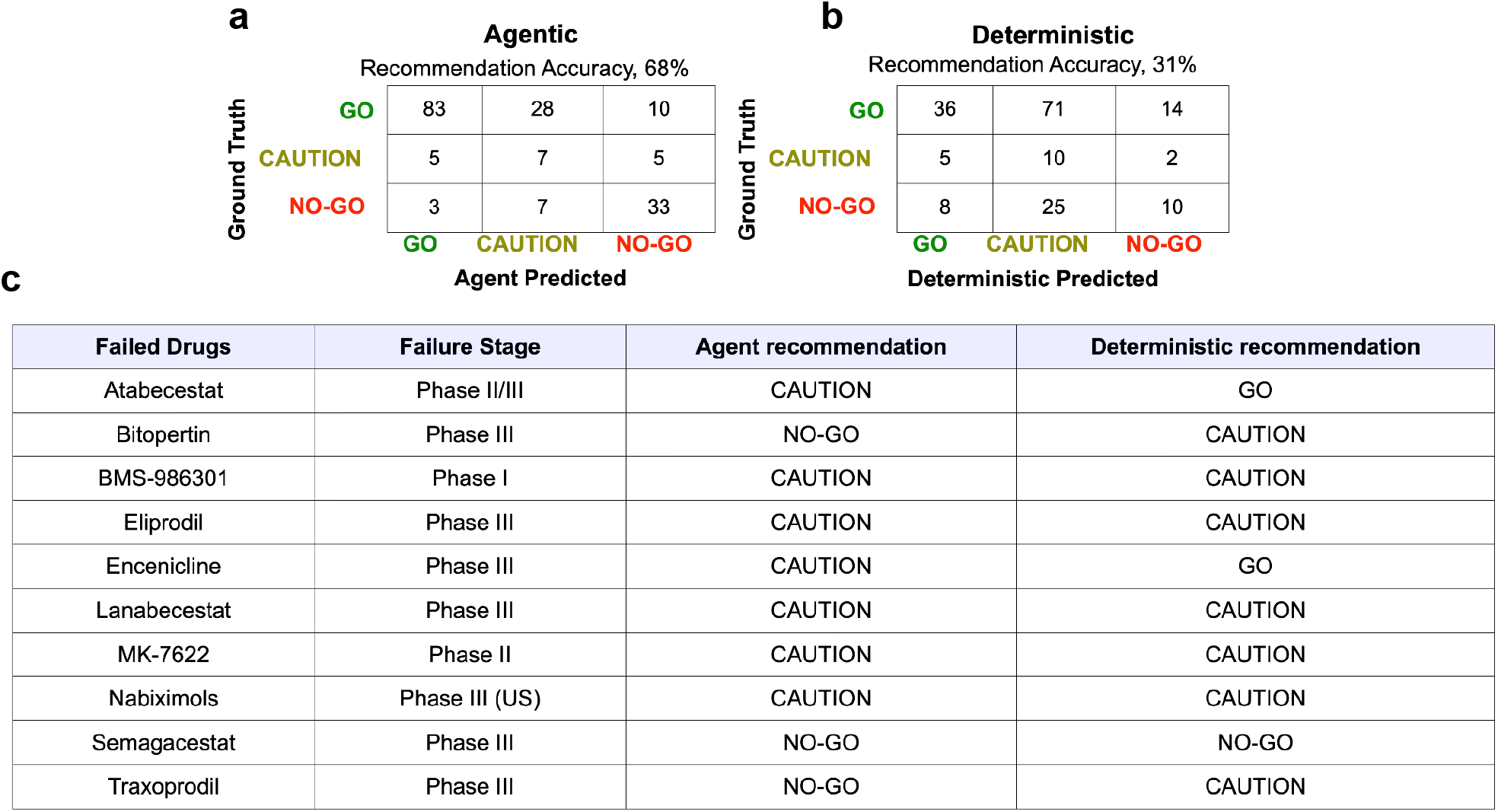
Comparative performance in identifying viable and failed CNS compounds. The agentic framework demonstrates superior predictive accuracy over the deterministic pipeline across n = 181 CNS compounds. **(a)** The Agentic framework achieved an overall recommendation accuracy of 68%. **(b)** The Deterministic pipeline achieved an accuracy of 31%. **(c)** Comparative accuracy analysis for ten representative failed CNS compounds, highlighting the discrepancy between the two approaches.

Confusion matrices reveal distinct error profiles for the two systems **(Figs. 3a, 3b)**. The agent correctly classified 83 of 121 GO compounds, 7 of 17 CAUTION compounds, and 33 of 43 NO-GO compounds. The agent’s primary error mode was over-caution, 28 GO compounds were classified as CAUTION. By contrast, the deterministic pipeline systematically over-assigned CAUTION labels, correctly classifying only 36 of 121 GO compounds while labeling 71 GO compounds as CAUTION falling below the majority-class baseline (66.9%) because its fixed thresholds are conservatively calibrated to pharmacological norms rather than tuned to this dataset. This pattern reflects the pipeline’s reliance on generic thresholds that trigger CAUTION flags without the pharmacological context needed to resolve them.

As a retrospective clinical failure detection system, the agent was evaluated on 16 clinically failed CNS candidates (one antibody, aducanumab, was excluded). Detection sensitivity varied by input tier, at Tier 1 (structure-only), the agent correctly flagged 87.5% (14/16) of clinical failures; at Tier 2 (with preclinical PK), 100% (9/9); and at Tier 3 (with measured Kp,brain), 100% (7/7). Detection was strongest for compounds that failed due to combined efficacy and safety signals, regulatory rejection, or cholinergic adverse events. Detection was weakest for compounds that failed due to lack of efficacy alone or gastrointestinal toxicity failure modes that lie outside the system’s pharmacokinetic and target-engagement modeling scope. Recommendation by Agent and deterministic pipelines for 10 representative failed drugs are shown in **Fig. 3c**.

### Eight pharmacology tools anchor the agent’s reasoning

The agent’s recommendation accuracy depends on the underlying mechanistic engine producing predictions that are directionally correct and appropriately ranked, even if point estimates carry substantial error. Allometric PK scaling from mouse to human achieved R^2^ = 0.76-0.92 across four parameters (CL, Vd, half-life, bioavailability), with 75-100% of predictions within 3-fold of observed values **(Fig. 2a)**. LogBB prediction achieved MAE = 0.39 log units (n = 83), with 69.9% of predictions within 0.5 log units of published values **(Fig. 2b)**, and Kp,brain estimates showed GMFE = 2.21 with 80% within 3-fold **(Fig. 2c)**. Receptor occupancy predictions, while carrying substantial absolute error (MAE = 31.5%, n = 73 dose-occupancy pairs across 14 targets), preserved dose-response monotonicity in all 23 multi-dose compounds **(Fig. 2d)**, the directional relationships essential for the agent’s contextual reasoning. Safety and transporter predictions showed balanced accuracies of 0.56–0.86 across individual tools **(Fig. 2e)**. PK conformal prediction intervals achieved 92.2% empirical coverage at 90% nominal level across 102 compounds **(Fig. 2f)**, providing calibrated uncertainty estimates that the agent used to flag unreliable downstream predictions.

### Reasoning traces reveal adaptive pharmacological interpretation

To illustrate the agent’s reasons, we present a detailed trace for two failed CNS drugs Atabecestat (BACE 1 inhibitor) **(Fig. 4a)** and Semagacestat (a gamma-secretase inhibitor) assessed across three input tiers **(Fig. 4b)**. At Tier 1 (SMILES-only), the agent sequentially invoked eight tools; BBB prediction, efflux assessment, active transport detection, safety screening, PK parameter estimation, PK simulation, PD modeling, and dose calculation. Each tool output was color-coded by provenance: at Tier 1, all pharmacokinetic inputs are predicted; at higher tiers, measured values progressively replace predictions.

**Fig 4.**
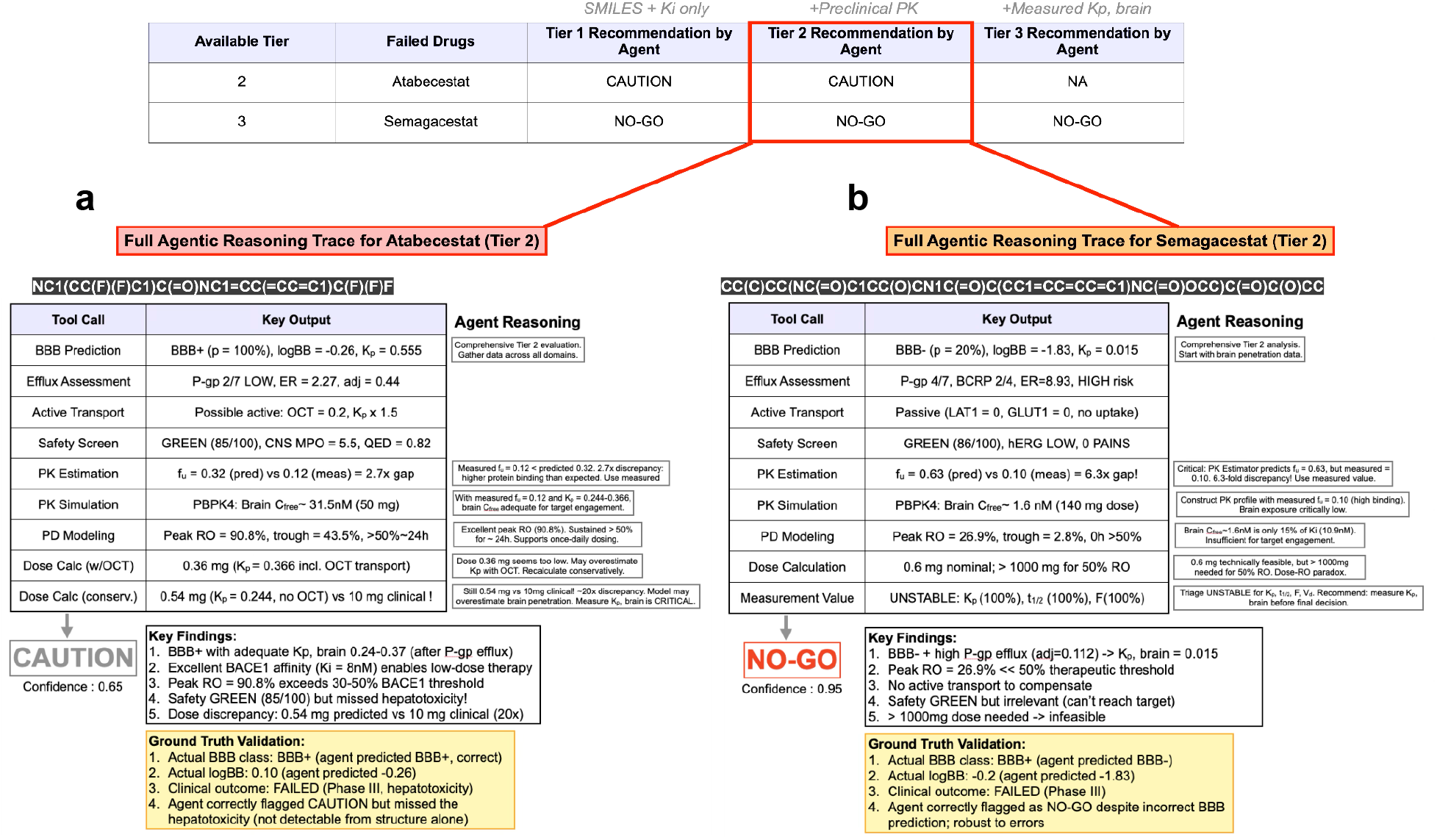
The Agentic framework performs reasoning traces and adaptive pharmacological interpretation within Tier 2 for failed CNS compounds. Based on these reasoning pathways, the agent recommended **(a)** CAUTION for Atabecestat. **(b)** NO-GO for Semagacestat. These traces reveal the underlying logic used to navigate complex pharmacological data that fixed rules might overlook.

The tier progression for these two failed drugs demonstrate the value of the tiered input system. Dose prediction fold-error decreased from 4.0 at Tier 1 to 1.2 at Tier 2 when measured mouse PK parameters replaced allometric estimates. Kp,brain fold-error similarly decreased from 10.3 at Tier 1 to 1.0 at Tier 3 when the measured brain-to-plasma ratio was provided. Across all compounds evaluated at multiple tiers, the fraction of measured (versus predicted) inputs increased from 0% at Tier 1 to 67% at Tier 2 and 83% at Tier 3, confirming that the tiered system operates as designed, each additional data tier replaces the least reliable predictions with measurements.

### Pharmacological reasoning outperforms fixed rules on discordant cases

To quantify the reasoning advantage, we compared recommendations on the 103 compounds (56.9% of the evaluation set) where the agent and deterministic pipeline disagreed **(Fig. 5a)**. Among these divergent cases, the agent was correct in 75% (77/103), the pipeline in 10% (10/103), and neither in 15.5% (16/103) (p = 1.48 × 10^-12, two-tailed exact binomial test on the 87 resolved cases). This result demonstrates that when the two systems disagree, the agent’s pharmacological reasoning is overwhelmingly more likely to produce the correct assessment.

**Fig 5.**
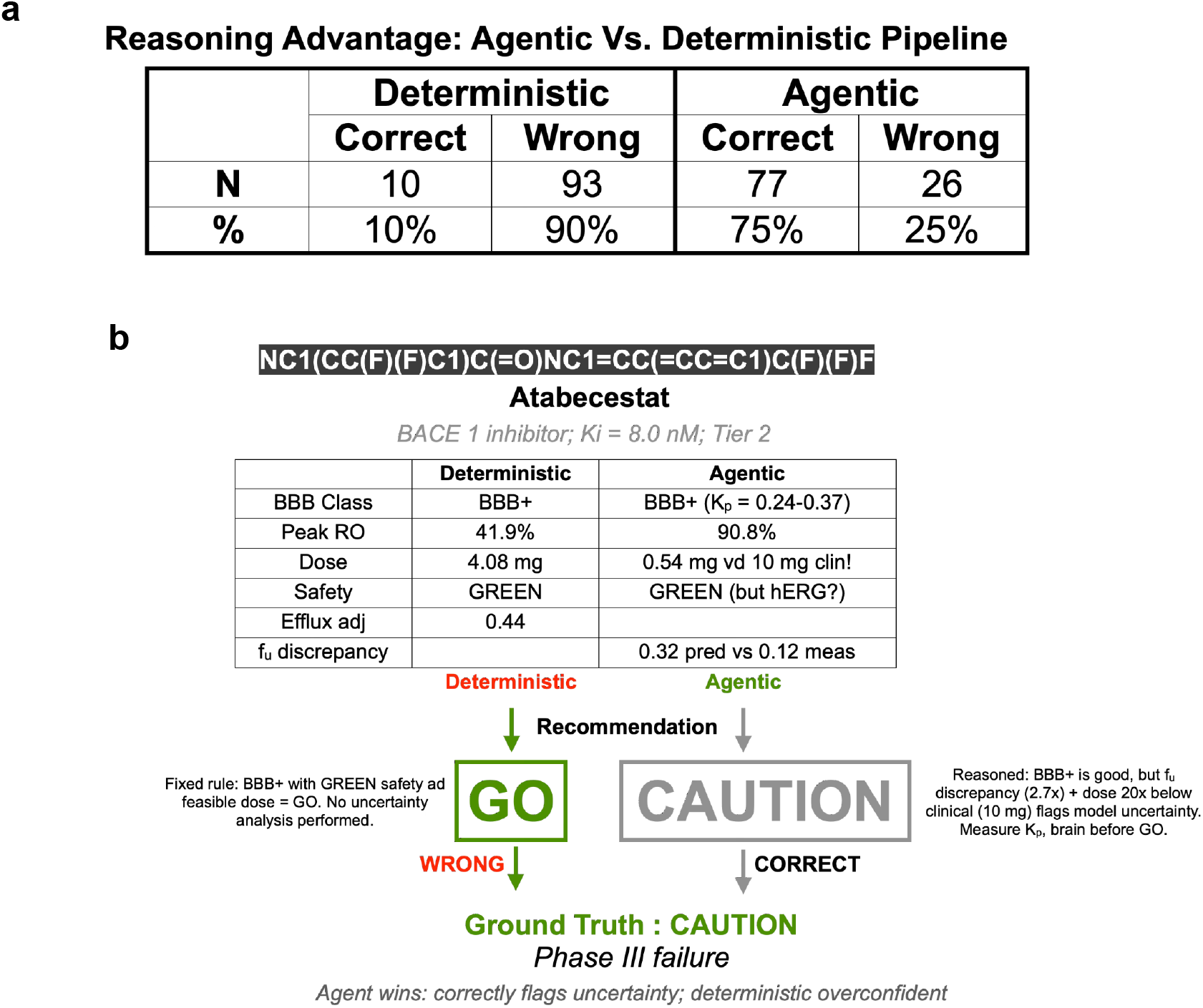
Superiority of pharmacological reasoning over fixed rules in discordant cases. Analysis of cases where the two architectures disagreed (discordant cases) highlights the robustness of the agentic approach.**(a)** In divergence analysis for the cases of conflicting recommendations, the Agentic framework was correct 75% of the time, compared to 10% for the deterministic pipeline.**(b)** A case study of Atabecestat serves as a representative example where the agent’s reasoning successfully outperformed the fixed-rule deterministic model.

The agent’s advantage was concentrated in specific drug classes. Among divergent cases, the agent outperformed the pipeline for atypical antipsychotics (8 agent wins), anticonvulsants (6 agent wins), SSRIs (6 agent wins), benzodiazepines (5 agent wins), SNRIs (4 agent wins), and TCAs (4 agent wins). The pipeline retained a modest advantage only for opioid analgesics (2 pipeline wins versus 1 agent win), a class where fixed occupancy thresholds are closer to pharmacological reality.

To illustrate how reasoning diverges from fixed rules, we present the case of the two failed CNS drugs **(Figs. 5b, 5c)**. The deterministic pipeline applied a fixed occupancy threshold and recommended GO for Atabecestat. The agent, recognizing the applied pharmacological reasoning about the target-specific occupancy requirements and recommended CAUTION, matching the ground truth **(Fig. 5b)**. This example typifies the pattern observed across divergent cases, the deterministic pipeline’s errors are not integration failures but reasoning failures. For Semagacestat, the pipeline and the agent recommended NO-GO but the agent provided deeper mechanistic justification, **Fig. 5c**.

Analysis of tool selection patterns revealed that the agent adapts its reasoning workflow to each compound’s pharmacological context. Across 181 compounds in agentic mode, the agent called a mean of 8.5 tools per compound (range: 0-18). The dose calculation tool was invoked for 78.5% of compounds, with the agent skipping dose estimation in 21.5% of cases typically when upstream predictions (e.g., BBB failure or high efflux) rendered dose estimation uninformative. The PD modeling tool was similarly skipped for 24.9% of compounds. This adaptive behavior contrasts with the deterministic pipeline, which executes the same fixed tool sequence for every compound. The overall evaluation reports for each 181 compounds using agentic and deterministic are available in **File 1** and **File 2** respectively.

## Discussion

The central finding of this work is that pharmacological reasoning over mechanistic model outputs produces measurably better compound assessments than the same models combined with fixed decision rules. When the agent and deterministic pipeline disagreed which occurred for more than half of all compounds the agent was overwhelmingly more likely to match the ground-truth recommendation. Both systems used identical mechanistic tools. The difference was entirely in how their outputs were synthesized. This supports the hypothesis that the bottleneck in computational preclinical CNS assessment is not model accuracy, but the integration step, deciding what a collection of interdependent predictions means for a specific compound at a specific target.

The per-class pattern of this advantage is informative. The agent outperformed the pipeline most strongly for drug classes where target-specific pharmacological knowledge overrides generic decision rules. GABA-A modulators (where the benzodiazepine binding site requires only 15-40% occupancy for anxiolysis), partial D2 agonists (where lower occupancy is therapeutic by design), and melatonergic agents (where standard dopaminergic thresholds do not apply). The pipeline retained a modest advantage for opioid analgesics, where the standard occupancy-efficacy relationship holds and the agent’s attempts at nuanced interpretation occasionally introduced errors. This asymmetry suggests that the reasoning advantage is a specific consequence of pharmacological knowledge being relevant and actionable. The same classes where a human pharmacologist would be expected to outperform a fixed rule, though we have not formally tested this comparison.

A recurring concern with LLM applications in drug discovery is that models may generate plausible but ungrounded assessments [26,27]. The architecture of *Cirrina* constrains this risk. The agent cannot invent pharmacokinetic data. Every number it reasons over is produced by a mechanistic tool with defined inputs, assumptions, and error characteristics. The agent’s role is to interpret, contextualize, and integrate these structured outputs not to generate them. The agent does not predict that a compound has 25% D2 occupancy; the Hill equation predicts this, and the agent interprets whether 25% is adequate given the compound’s target pharmacology. The mechanistic engine’s accuracy was validated to be sufficient for this interpretive role, though not for standalone quantitative prediction. Notably, receptor occupancy predictions showed correct dose-response monotonicity in all multi-dose compounds, preserving the directional relationships essential for reasoning despite substantial point-estimate error. The agent’s ability to flag downstream predictions as unreliable when upstream inputs are uncertain skipping receptor occupancy estimation when BBB penetration is predicted to be negligible, and represents a form of epistemic awareness that fixed pipelines lack.

*Cirrina* occupies a position between two established paradigms in computational pharmacology. Free ADMET predictors (pkCSM, SwissADME, ADMETlab) provide endpoint-level classification with reported BBB accuracies of 85-92% [10-12], but produce isolated predictions without mechanistic chaining, uncertainty quantification, or integrative reasoning. Enterprise PBPK platforms (Simcyp, GastroPlus) model the full mechanistic chain with higher fidelity, but require extensive measured inputs unavailable during early lead optimization and do not incorporate adaptive reasoning. *Cirrina* bridges this gap. It operates from minimal data (structure-only at Tier 1), chains predictions mechanistically, and reasons over the chain to produce an integrated assessment.

The distinction from recent LLM applications in drug discovery [21-23,28,29] is architectural. Those approaches use LLMs primarily as prediction engines, mapping inputs to outputs through learned representations, whereas *Cirrina* uses the LLM as a reasoning engine over externally computed mechanistic outputs closer to tool-augmented reasoning [19,20] than to end-to-end prediction. This trades raw prediction performance for interpretive transparency. Every intermediate result is traceable to a specific mechanistic tool, and the agent’s reasoning is inspectable through its tool-call sequence and natural-language rationale. For pharmaceutical applications, where regulatory and institutional requirements favour interpretability, this transparency may matter as much as accuracy.

Integration into drug discovery workflows would likely occur at two stages. During lead optimization, the system could provide rapid first-pass assessments of compound series, flagging candidates with unfavourable pharmacological profiles before resource-intensive in vivo studies. During candidate selection, the tiered input system allows progressive refinement as preclinical data accumulates the same compound can be re-evaluated at higher tiers as PK, brain penetration, and safety data become available, with each tier replacing the least reliable predictions with measurements.

The choice of LLM warrants discussion. We used Claude Haiku 4.5, a relatively small model optimized for speed and cost, rather than a larger frontier model. This was a deliberate trade-off, at approximately 181 compounds per evaluation run, the cost and latency of frontier models would be prohibitive for iterative development. Whether larger models with deeper reasoning capabilities would improve recommendation accuracy particularly for the 15.5% of divergent cases where neither system was correct remains an open question. The system prompt, which encodes pharmacological knowledge and reasoning principles rather than prescriptive rules, is model-agnostic. The architecture could be evaluated with different LLMs without modification to the tools or prompting strategy. Such a comparison would help disentangle the contributions of the reasoning architecture from the specific model’s capabilities. We used temperature = 0 to minimize output variability. Characterizing inter-run reproducibility and prompt sensitivity [30] across the ∼554-line system prompt would further strengthen confidence in the architecture and are natural next steps for translational deployment.

Several directions for future work would strengthen and extend these findings. First, and most importantly, prospective validation on proprietary compounds would address the concern that the LLM’s training data includes pharmacological information about the 181 known compounds evaluated here. Leave-one-class-out cross-validation provides partial mitigation, seven drug classes achieve F1 = 1.0, suggesting the advantage is not driven by class-level memorization but a blinded evaluation on compounds from an active drug discovery programme would provide the definitive test, alongside expanding the evaluation set to improve statistical power for the CAUTION class. Currently 17 compounds; we address the current class imbalance of 121 GO, 17 CAUTION, 43 NO-GO through MCC, macro F1, and majority-class baseline comparison, but a more balanced dataset would strengthen conclusions. Second, in the model level, correcting the systematic overprediction in receptor occupancy through empirical calibration against PET data, tissue-binding corrections, or non-equilibrium receptor kinetics models would improve the inputs available to the agent’s reasoning. Third, extending conformal calibration across the full prediction chain would close the gap between PK-level coverage (92.2% at 90% nominal) and end-to-end occupancy-level coverage (33.6%). The current gap reflects uncalibrated error from receptor binding parameters and model structure that PK intervals cannot absorb, and counterintuitively Tier 1 coverage exceeds Tier 2 because wider prediction intervals produce wider occupancy bounds while measured PK data narrows the intervals without reducing pharmacodynamic error. A Bayesian hierarchical approach that jointly models PK and PD uncertainty, or direct conformal calibration of model outputs against clinical data, could address this. Fourth, incorporating organ-specific toxicity predictions and disease-model efficacy endpoints would extend clinical failure detection. Currently strong for safety-signal compounds (87.5% at Tier 1, 100% at higher tiers) into failure modes such as gastrointestinal toxicity and isolated efficacy failure that lie outside the current modelling scope. Though predicting therapeutic inadequacy from preclinical pharmacology alone remains a fundamental challenge. Finally, the reasoning-over-mechanistic-models paradigm may generalize beyond CNS to therapeutic areas with analogous interdependencies such as oncology (target accessibility, tumour penetration, resistance) or immunology (target-mediated drug disposition), though the specific tools and pharmacological knowledge would need to be rebuilt for each domain.

*Cirrina* for the first time demonstrates that combining mechanistic pharmacological models with LLM-driven reasoning produces compound assessments that exceed the performance of either component alone. The system addresses the integration gap, the manual, undocumented step where interdependent predictions are synthesized into a pharmacological assessment. By automating this synthesis through a reasoning loop that is inspectable and reproducible in architecture. The mechanistic engine provides grounded, traceable predictions. The LLM agent provides pharmacological interpretation that adapts to target context, upstream uncertainty, and drug-class-specific knowledge. The conformal prediction framework provides calibrated uncertainty at the PK level with a clear path toward end-to-end calibration. The result is not a replacement for experimental pharmacology or clinical judgement, but a tool that can provide a structured, mechanistically grounded first assessment of any CNS compound from its structure alone, an assessment that would otherwise require expert pharmacological analysis.

